# Chaperone-mediated heterotypic phase separation prevents the amyloid formation of the pathological Y145Stop variant of the prion protein

**DOI:** 10.1101/2024.09.05.611572

**Authors:** Lisha Arora, Dipankar Bhowmik, Snehasis Sarkar, Anusha Sarbahi, Sandeep K. Rai, Samrat Mukhopadhyay

**Author notes:** Corresponding author: Samrat Mukhopadhyay. Contributed equally (Lisha Arora, Dipankar Bhowmik). Department of Stem Cell and Regenerative Biology, Harvard University, Cambridge, USA.

## Abstract

Biomolecular condensates formed via phase separation of proteins and nucleic acids are crucial for the spatiotemporal regulation of a diverse array of essential cellular functions and the maintenance of cellular homeostasis. However, aberrant liquid-to-solid phase transitions of such condensates are associated with several fatal human diseases. Such dynamic membraneless compartments can contain a range of molecular chaperones that can regulate the phase behavior of proteins involved in the formation of these biological condensates. Here, we show that a heat shock protein 40 (Hsp40), Ydj1, exhibits a holdase activity by potentiating the phase separation of a disease-associated stop codon mutant of the prion protein (Y145Stop) either by recruitment into Y145Stop condensates or via Y145Stop-Ydj1 two-component heterotypic phase separation that prevents the conformational conversion of Y145Stop into amyloid fibrils. Utilizing site-directed mutagenesis, multicolor fluorescence imaging, single-droplet steady-state and picosecond time-resolved fluorescence anisotropy, fluorescence recovery after photobleaching, and fluorescence correlation spectroscopy, we delineate the complex network of interactions that govern the heterotypic phase separation of Y145Stop and Ydj1. We also show that the properties of such heterotypic condensates can further be tuned by RNA that promotes the formation of multicomponent multiphasic protein-RNA condensates. Our vibrational Raman spectroscopy results in conjunction with atomic force microscopy imaging reveal that Ydj1 effectively redirects the self-assembly of Y145Stop towards a dynamically-arrested non-amyloidogenic pathway, preventing the formation of typical amyloid fibrils. Our findings underscore the importance of chaperone-mediated heterotypic phase separation in regulating aberrant phase transitions and amyloid formation associated with a wide range of deadly neurodegenerative diseases.

## Introduction

Eukaryotic cells organize their complex array of biochemical processes within the crowded cellular milieu by compartmentalizing biomolecules such as proteins, lipids, and nucleic acids into the well-known membrane-enclosed organelles such as the Golgi complex, nucleus, endoplasmic reticulum, and so forth. Studies in the past decade and a half have revealed that, in addition to these conventional organelles, cells also contain noncanonical membraneless organelles known to form via phase separation of proteins, nucleic acids, and other biomolecules. These membraneless compartments, also known as biomolecular condensates, can participate in a myriad of cellular processes, including DNA repair, signaling, transcription, ribosome biogenesis, and protein quality control (PQC) (1–10). A growing body of research has shown that intrinsically disordered proteins/regions (IDPs/IDRs) often comprising prion-like, low-complexity domains are the key candidates for biological phase separation due to their conformational flexibility, structural heterogeneity, and their ability to engage in a multitude of transient, non-covalent interactions, such as electrostatic, hydrophobic, cation-π, and π-π interactions. These interactions result in the formation of viscoelastic complex fluids involving density transition coupled with percolation and coupled associative and segregative phase transitions (11–22). Such phase-separated condensates are highly dynamic assemblies that can form and dissolve in response to cellular cues. However, abnormal phase transitions of such dynamic liquid-like condensates into solid-like protein aggregates have been associated with a range of human diseases (23–26). An intriguing example is the mammalian prion protein (PrP) which is proposed to follow two distinct pathways to infectious protein aggregates namely, via the conventional protein misfolding pathway (27, 28) and the phase separation-mediated pathway (29).

The conformational conversion of the cellular form of the prion protein (PrP^C^) into a β-rich, aggregation-prone, autocatalytic, self-templating scrapie form (PrP^Sc^) that is associated with a range of fatal transmissible neurogenerative diseases collectively known as prion diseases. PrP^C^ is a 253-residue glycosylphosphatidylinositol-anchored (GPI-anchored) cell-surface protein consisting of an N-terminal signal peptide (residues 1-23) that is cleaved upon maturation, a positively charged intrinsically disordered N-terminal tail (residues 23-120), a C-terminal globular domain (residues 121-231), and a GPI-anchor signal (residues 231-253) (27–31). The destabilization of the folded globular domain of PrP^C^ is thought to trigger a cascade of misfolding and aggregation events resulting in the formation of amyloid-like infectious protein aggregates that are capable of binding and autocatalytically converting PrP^C^ into PrP^Sc^ (32). In contrast, the N-terminal intrinsically disordered region (IDR) possessing five octapeptide repeats that is thought to be functionally important for copper binding can promote phase separation of PrP. The N-terminal IDR is proposed to promote phase separation, whereas the C-terminal globular domain can potentially regulate phase separation (29, 33, 34). The phase behavior of the PrP is influenced by cofactors like RNA, DNA, and metal ions that tune the material properties of these condensates (33–38). A naturally occurring amber stop codon mutant (Y145XStop, also represented as Y145X) results in a C-terminally truncated, intrinsically disordered variant of the protein that is associated with a Gerstmann-Sträussler-Scheinker (GSS)-like phenotype and familial cerebral amyloid angiopathy (Fig. 1A) (39–44). Under normal conditions, Y145X is rapidly degraded by the proteasome-mediated pathway; however, under stress conditions, it gets mislocalized and accumulates in the nucleus and cytosol (45). We demonstrated that this intrinsically disordered Y145X spontaneously phase separates into highly dynamic, liquid-like droplets that readily mature into ordered, solid-like, self-templating amyloid aggregates via liquid-to-solid phase transitions (33). These studies hinted at a protective role of the C-terminal globular domain in regulating aberrant phase transitions of the N-terminal IDR of PrP.

**Fig. 1.**
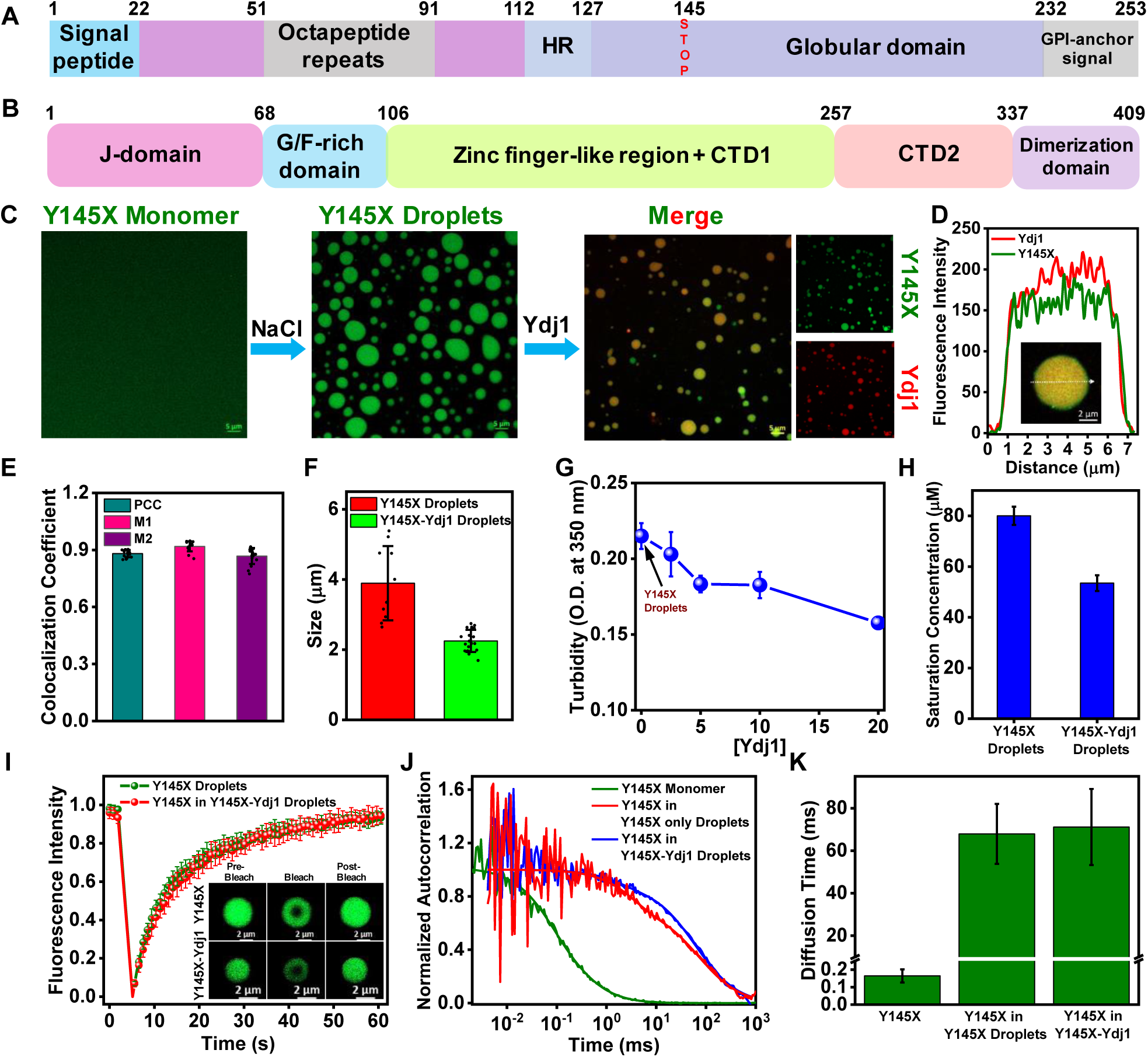
Recruitment of Ydj1 within Y145X condensates. (A) Domain architecture of full-length human PrP showing all the segments (Y145X is highlighted). (B) Schematic representation of Ydj1 indicating all the domains. (C) A high-resolution confocal image of a dispersed phase and droplets of Y145X containing 0.1% AlexaFluor488-labeled Y145X single-Cys mutant at residue 31 (W31C) (100 μM protein, 20 mM sodium phosphate, 350 mM NaCl, pH 7.5, scale bar 5 μm), two-color Airyscan confocal images of colocalized Y145X (W31C Y145X-AlexaFluor488, green) and Ydj1 (labeled with AlexaFluor594, red) in Y145X-Ydj1 condensates (scale bar 5 μm). (D) Fluorescence intensity profiles showing the colocalization of Y145X and Ydj1 (inset shows the representative colocalized image of the droplet) (E) Pearson’s correlation coefficient (PCC) and Manders’ overlap coefficients plots quantifying the fraction of Ydj1 colocalizing with Y145X (M1) and fraction of Y145X colocalizing with Ydj1 (M2) (The data represent mean ± SD; n ≥ 15 high-resolution confocal images from five independent reactions). (F) Bar plot of the size of the droplets formed by Y145X in the absence and presence of Ydj1 (The data represent mean ± SD; n ≥ 10 high-resolution confocal images from five independent reactions). (G) Turbidity measurements of Y145X-Ydj1 reaction mixtures as a function of increasing Ydj1 concentration. The concentration of Y145X was kept constant at 100 μM (350 mM NaCl, 20 mM sodium phosphate, pH 7.5). The data represent mean ± SD; n = 3. (H) Saturation concentration (C_sat_) of Y145X for Y145X-only and Y145X-Ydj1 condensates estimated by sedimentation assay. (I) FRAP kinetics of Y145X for Y145X only droplets and Y145X-Ydj1 droplets using AlexaFluor488-labeled Y145X (n = 10). (J) Normalized autocorrelation plots obtained from FCS measurements performed for monomeric Y145X and for Y145X in Y145X only droplets and Y145X-Ydj1 droplets (unnormalized autocorrelation plots are shown in Fig. S1). (K) Diffusion time of Y145X in monomeric dispersed form, in Y145X only droplets and in Y145X-Ydj1 droplets estimated from FCS measurements (Data represent mean ± SD for n = 5 independent samples). For FCS measurements, 5 nM of W99C Y145X-AlexaFluor488-labeled protein was used for the monomeric dispersed phase, 0.25 nM of AlexaFluor488-labeled Y145X (W99C) was used for droplet measurements. More details are provided in the supporting information.

Despite extensive research, it remains unclear how cellular homeostasis is maintained and the formation of toxic protein aggregates is prevented. A growing body of evidence highlights the multifaceted role of molecular chaperones in the cellular PQC system in reducing pathogenesis and maintaining protein homeostasis within cells (46, 47). Chaperones assist client proteins in folding and translocation and maintain clients in their native conformation, avoiding misfolding and aggregation. Proteomic analysis of stress granules has revealed the presence of a diverse array of molecular chaperones, including heat shock proteins (Hsps) such as Hsp40, Hsp70, Hsp90, and small Hsps, underscoring the importance of chaperones in the formation and maintenance of membrane-less organelles (48–51). Of all, Hsp40 is a versatile protein that functions as a co-chaperone in the Hsp70 machinery, responsible for both recognizing and targeting clients to the chaperones and activating the Hsp70 chaperone cycle (52–55). Proteins of the Hsp40 family possess a highly conserved domain architecture across different species comprising of an N-terminal J-domain, a glycine-phenylalanine (G/F) linker, a zinc-finger-like region, and peptide binding C-terminal domains (CTD I/II), followed by a dimerization-domain (Fig. 1B) (56, 57). Recent studies have shown that Hsp40 not only assists Hsp70 in its chaperone functions but also exhibits holdase activity independently, thereby preventing abnormal protein aggregation (58). In this work, we demonstrate that Ydj1, a yeast homolog of the human class-I Hsp40, Hdj2 (DNAJA1), which is known to localize in the nucleoli under normal conditions, also gets relocated to the stress granules in the cytoplasm during stress response, modulates the phase behavior Y145X. Furthermore, we also show that Ydj1 can regulate the physical properties of the phase-separated condensates formed in the presence of cofactors like RNA and can prevent aberrant phase transition and aggregation of Y145X. Such a phase separation-mediated chaperoning of Y145X by Ydj1 underscores the critical role of the Hsp40 chaperone in preserving the dynamic native state of proteins and in preventing pathological protein aggregation.

## Results

### Ydj1 gets recruited within phase-separated droplets of prion Y145X

Y145X is intrinsically disordered with a high net positive charge (+10.4) at pH 7.5 and contains a weakly hydrophobic region (HR; residues 113 to 135) enabling it to undergo phase separation via a range of multivalent interactions as previously demonstrated by us (Fig. S1A) (33). First, to establish the structural similarity between yeast Hsp40 and human Hsp40, we compared the structure of Ydj1 and human analog Hdj2 (DNAJA1). Structural superimposition revealed a striking similarity in their three-dimensional structures. This similarity is supported by other studies showcasing the functional robustness across various species, highlighting their evolutionary conservation (59). The sequence analysis of Ydj1 using FuzDrop, along with confocal microscopy imaging of Ydj1 (10 µM) doped with 1% AlexaFluor594-C5-maleimide labeled Ydj1, indicated that Ydj1 did not phase separate under our experimental conditions (Fig. S1B, C). We then began by inducing the phase separation of Y145X (100 µM Y145X, 350 mM NaCl, 20 mM sodium phosphate, pH 7.5). Next, we set out to see the effect of Ydj1 on these pre-formed phase-separated condensates of Y145X. Our results demonstrated the recruitment of Ydj1 and its colocalization within Y145X droplets, which was further corroborated by the Pearsons’ coefficient colocalization and Manders’ overlap analysis plots (Fig. 1C-E). However, further analysis showed a decrease in the size of Y145X droplets upon the introduction of Ydj1 and a concentration-dependent reduction in the turbidity of Y145X droplets with increasing Ydj1 concentrations, suggesting that Ydj1 significantly influences the phase behavior of Y145X (Fig. 1F, G, S1D). Next, we estimated the saturation concentration (C_sat_) of Y145X for Y145X only and Y145X-Ydj1 droplets and found it to be lower in the presence of Ydj1 (Fig. 1H). We then performed fluorescence recovery after photobleaching (FRAP) experiments, which showed a near-complete recovery for the Y145X in the absence and presence of Ydj1, suggesting that the droplets retained their liquid-like characteristics even in the presence of Ydj1 (Fig. 1I). To further investigate the diffusion within the individual droplets formed on a microsecond to sub-millisecond timescale, we monitored the diffusion time of AlexaFluor488-labeled Y145X (W99C) inside Y145X-only and Y145X-Ydj1 droplets using fluorescence correlation spectroscopy (FCS). The diffusion time extracted from our correlation measurements suggested slower diffusion of Y145X within the droplets formed in the absence and presence of Ydj1, suggesting the formation of a densely interconnected network inside these condensates (Fig. 1J, K, Fig. S1E-G). Taken together, our results demonstrate that the recruitment of Ydj1 regulates the phase separation of Y145X and results in the formation of a stronger viscoelastic network within the droplets constituted by the two proteins. Having established the recruitment of Ydj1 in Y145X condensates, we next aimed to discern the intricate network of interactions that govern the co-partitioning of Ydj1 into Y145X droplets.

### An interplay of hydrophobic and electrostatic interactions governs the recruitment of Ydj1 in Y145X droplets

Hsp40 is known to interact through its polypeptide binding domain (PBD), which recognizes and interacts with the hydrophobic side chains of non-native or misfolded proteins, thereby preventing them from aggregation (60). Therefore, we started by discerning the role of hydrophobic interactions in driving the recruitment of Ydj1 into Y145X condensates. We first analyzed the hydropathy plot and amino acid sequence of Y145X, which revealed the presence of a hydrophobic region (HR) rich in alanine and valine (Fig. 2A, S2A). Since valine is more hydrophobic than alanine, we mutated three alanine-to-valines (A→V Y145X) at residue positions 113, 115, and 117 to increase the hydrophobicity of this region. As shown previously, Y145X phase separation is also driven by hydrophobic interactions; the mutation resulted in more gel-like droplets with a very low fluorescence recovery (FRAP) (Fig. S2B) (33). However, when Ydj1 is introduced into the A→V Y145X droplets, a complete colocalization was observed, as can be seen in two-color confocal microscopy imaging, along with an increase in the FRAP recovery of Y145X, which is further supported by the half-time plot (Fig. 2B-D, S2C). These findings suggest that Ydj1 indeed interacts with the hydrophobic region of wild-type Y145X. At physiological pH, Y145X exists as a positively charged polypeptide (pI = 11.6), while Ydj1 exhibits a negative charge under the same condition (pI = 6.30) (61). Therefore, we next postulated that electrostatic interactions might also facilitate the interaction of Ydj1 into Y145X droplets at physiological pH. To experimentally verify this, we carefully looked at the sequence and net charge per residue plot of Y145X and created single cysteine (Cys) mutants at three positions: 31 (near a highly positively charged cluster); 99 (near a moderately charged cluster); 120 (with no charged cluster) (Fig. 2E, S2D). We then employed site-specific single-droplet steady-state fluorescence anisotropy measurements to delineate the regio-specific interactions between Y145X and the chaperone upon the addition of Ydj1 in the preformed Y145X droplets (62). Fluorescence anisotropy measurements report the local rotational flexibility of the fluorophore; an increase in the steady-state anisotropy is typically interpreted as a loss of conformational flexibility, indicating the formation of interactions. To record the site-specific anisotropy, we used single-Cys variants labeled with thiol-active fluorescein-5-maleimide (F5M). An increase in anisotropy was observed at all three residue positions compared to monomeric dispersed Y145X, indicating a substantial reduction in the conformational flexibility and rotational mobility of the Y145X polypeptide within the condensed phase. Interestingly, upon the addition of Ydj1 to preformed Y145X droplets, a further enhancement in anisotropy was observed at locations 31 and 99, indicating the involvement of electrostatic interactions between the positively charged clusters of the Y145X polypeptide and the negatively charged Ydj1 within the phase-separated droplets (Fig. 2F, G).

**Fig. 2.**
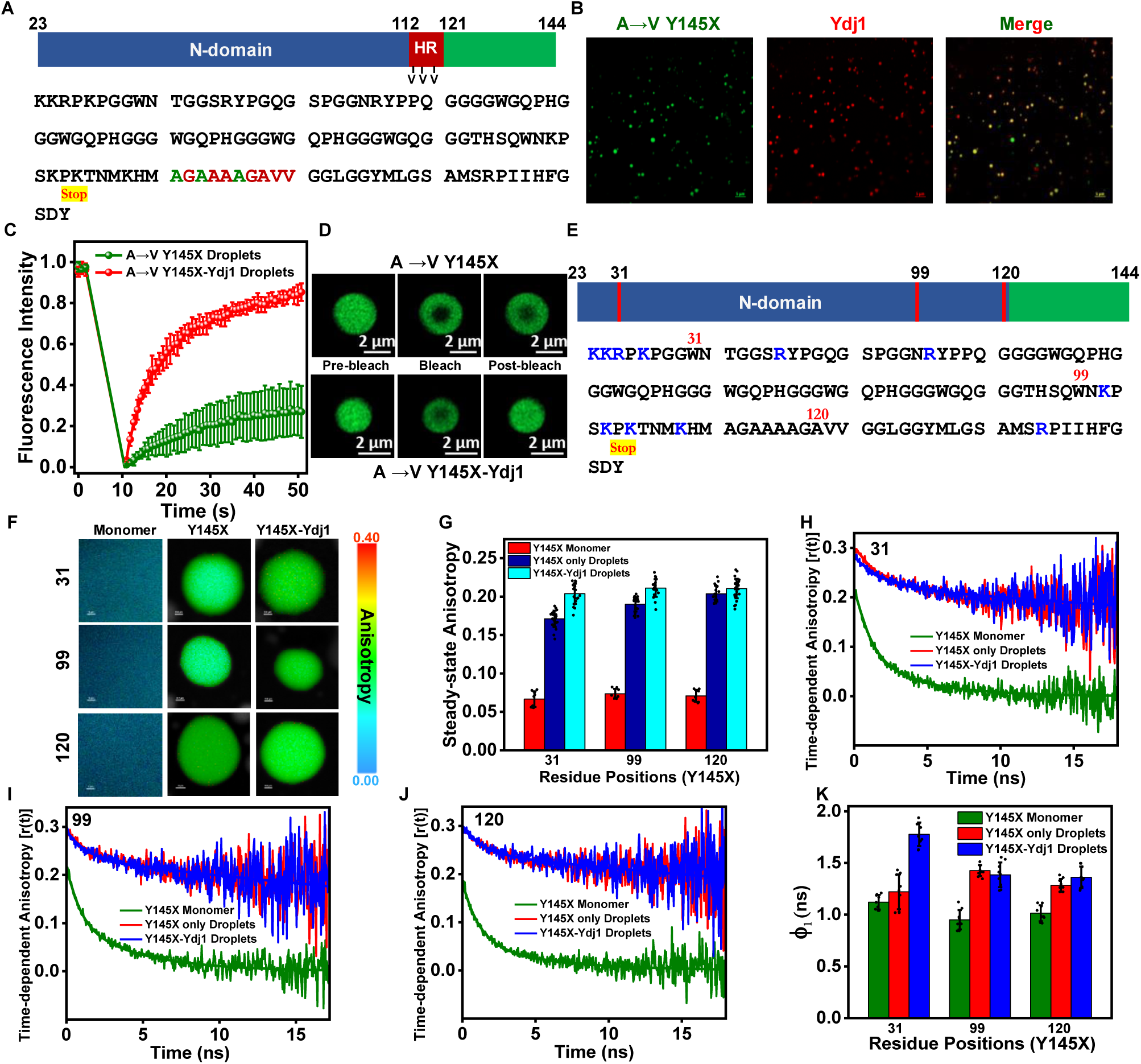
An interplay of hydrophobic and electrostatic interactions governs the recruitment of Ydj1 in Y145X droplets. (A) Schematic representation and amino acid sequence of Y145X highlighting the hydrophobic region (maroon) and the residue positions mutated to Valine (olive). (B) Two-color Airyscan confocal images of colocalized A→V Y145X variant (AlexaFluor488-labeled, green) and Ydj1 (AlexaFluor594-labeled, red) in Y145X-Ydj1 condensates (scale bar 5 μm) after the addition of Ydj1 in A→V Y145X droplets. (C) FRAP kinetics of A→V Y145X only droplets and droplets formed after addition of Ydj1 to the preformed droplets (The data represent mean ± SD; n ≥ 5). (D) Fluorescence images of droplets obtained during FRAP measurements (scale bar 2 µm). (E) Schematic representation and amino acid sequence of Y145X highlighting the single-Cys mutants (red). (F) Representative fluorescence anisotropy images of the monomeric dispersed phase and a single-droplet of single-Cys variants of Y145X labeled with Fluorescein-5-maleimide (F5M). (G) Steady-state fluorescence anisotropy of single-Cys Y145X labeled with F5M at different positions in the monomeric dispersed phase (red), in Y145X only droplets (navy), and in droplets after the addition of Ydj1 (cyan). Data represents mean ± SD of n ≥ 8 independent samples for monomers and n ≥ 20 for droplets. (H-J) Representative time-resolved fluorescence anisotropy decays of F5M-labeled Y145X at residue positions 31, 99, and 120 in monomeric dispersed phase (green), Y145X only droplets (red) and Y145X-Ydj1 droplets (blue). The solid lines are fits obtained using decay analysis. See *Supporting Information*, Table S2 for recovered parameters. (K) Plot showing the fast rotational correlation time (ϕ) of Y145X for different residue positions in the dispersed phase, Y145X only droplets, and Y145X-Ydj1 droplets. Data represents mean ± SD of n = 9.

Since the steady-state anisotropy provides time-averaged information, it cannot distinguish different modes of rotational motion of the polypeptide chain. Next, we utilized picosecond time-resolved fluorescence anisotropy measurements to unravel distinct modes of rotational dynamics occurring at the nanosecond time scale using fluorescently labeled single-Cys variants of Y145X. In the monomeric form, F5M-labeled Y145X exhibited fast depolarization kinetics for all three residue positions (residues 31, 99, and 120), characteristic of a rapidly fluctuating expanded IDP. The depolarization kinetics were described by a biexponential decay function comprising a (fast) local-rotational correlation time representing a local probe motion (∼ 1 ns) of the fluorophore and a slow rotational correlation time (∼ 4-5 ns) corresponding to backbone dihedral motions and long-range reorientation fluctuations. Upon homotypic phase separation, Y145X exhibited slower depolarization kinetics, indicating dampening of the chain rotational dynamics due to chain-chain interaction within the condensates. Interestingly, the addition of Ydj1 to the preformed Y145X condensates resulted in even slower depolarization kinetics at residue position 31 (Fig. 2H-J, Table S2). The fast sub-nanosecond rotational correlational time, which corresponds to the local rotational dynamics of the fluorophore, also showed an increase at position 31 likely due to a more constrained environment surrounding the positive charge cluster. This suggests the formation of electrostatic interactions, which supports our steady-state anisotropy findings. (Fig 2K). Taken together, these findings indicate an interplay of both hydrophobic and electrostatic interactions that drive the recruitment of Ydj1 in Y145X condensates. Since these condensates can promote aberrant liquid-to-solid phase transitions into amyloids, we next set out to elucidate the role of the Ydj1 in regulating liquid-to-solid phase transitions.

### Ydj1 inhibits liquid-to-solid phase transitions of Y145X droplets

Y145X droplets readily undergo liquid-to-solid phase transitions leading to the formation of amyloid-like aggregates as reported by us. (Fig. 3A) (33). Our time-dependent confocal imaging showed that in the presence of Ydj1, the Y145X droplets retained the phase-separated state when monitored for 12 hours and did not convert into solid-like aggregates (Fig. 3B). To gain deeper insights into the structural properties of the species formed on maturation, we utilized vibrational Raman spectroscopy and monitored the amide I vibrational band (1620 - 1720 cm^-1^). This band originates from the C=O stretching vibrations and is detected within the 1620-1720 cm^-1^ range and is a direct indicator of the hydrogen-bonded cross-β amyloid architecture. Our vibrational Raman spectroscopy analyses revealed a relatively broad amide I band for Y145X droplets and a sharp amide I peak at ∼ 1675 cm^-1^ after 12 hours of incubation indicating the time-dependent maturation of Y145X-only droplets into an ordered amyloid-like state as observed by us previously (Fig. 3C) (33). In contrast, the condensed phase of Y145X droplets formed after the recruitment of Ydj1 exhibited a broad amide I band even upon 12 hours of incubation indicating the inhibition of conformational conversion into a solid-like amyloid state (Fig. 3D). Since the aggregates formed upon aging of Y145X condensates transition into amyloid-like aggregates are thioflavin T (ThT) positive, we next wanted to test the impact of Ydj1 on the aggregation kinetics of Y145X droplets in a dose-dependent manner. To achieve this, we initiated the aggregation reaction of Y145X droplets using ThT as the amyloid reporter under a stirring condition. The phase-separation mediated aggregation of the Y145X followed a nucleation-dependent polymerization mechanism with a lag time of ⁓ 30 minutes. We then performed atomic force microscopy (AFM) of the species formed upon saturation of the reaction, and as expected, a fibrillar nanoscale species with a height of around 6 nm was observed. In contrast, the aggregation reaction performed in the presence of Ydj1 showed a decrease in the ThT intensity with its increasing concentration, thereby deaccelerating the aggregation, indicating the Ydj1-mediated abrogation of Y145X aggregation (Fig. 3E, F). The AFM imaging of the final species formed in the Y145X-Ydj1 reaction mixture revealed small, heterotypic protein clusters, which can be described as protein conglomerates (Fig. 3G, H). Taken together, our results reveal that Ydj1-mediated heterotypic phase separation of Y145X inhibits liquid-to-solid phase transitions and amyloid formation. The chaperoning activity of Ydj1 appears to redirect the self-assembly of Y145X into a non-amyloidogenic pathway, leading to the formation of heterogeneous non-amyloid protein clusters. Since Y145X is also known to undergo aggregation via a typical nucleation-dependent polymerization pathway, therefore, we next set out to delineate the role of Ydj1 on monomeric Y145X under a condition that promotes the aggregation of monomeric Y145X into pathological amyloid fibrils.

**Fig. 3.**
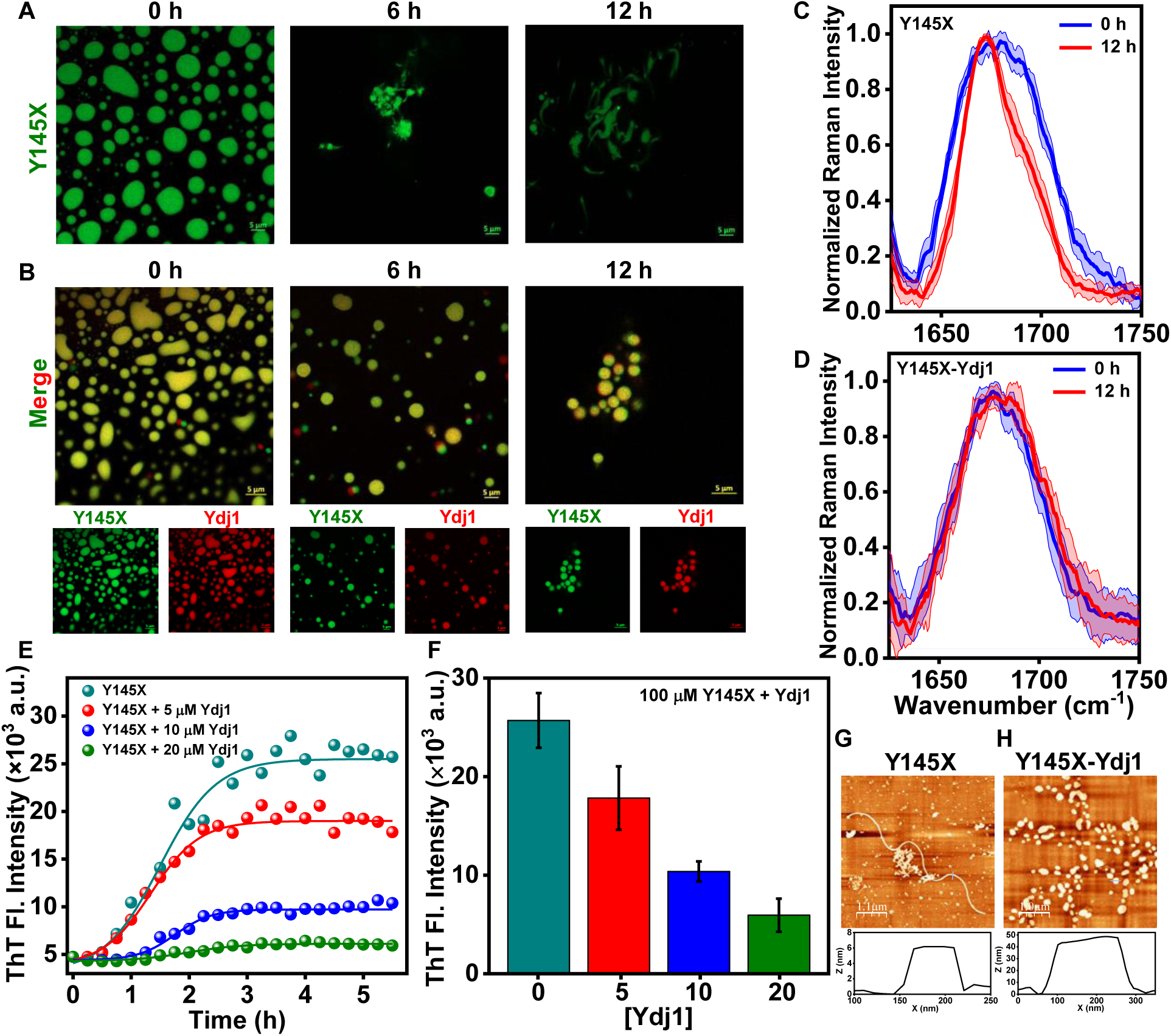
Ydj1 halts the aggregation of Y145X droplets. (A) Time-dependent high-resolution confocal images of Y145X-only droplets showing morphological transitions (using 0.1% AlexaFluor488-labeled Y145X, scale bar 5 μm). (B) Two-color Airyscan confocal images of colocalized Y145X (AlexaFluor488-labeled, green) and Ydj1 (AlexaFluor594-labeled, red) in Y145X-Ydj1 condensates (scale bar 5 μm) as a function of time. (C), (D) Amide I vibrational Raman band of the dense phase of Y145X and Y145X-Ydj1 recorded after 10 minutes of incubation and 12 hours of incubation. (E) Thioflavin T (ThT) kinetics of phase separation-mediated aggregation of Y145X-only droplets and with increasing concentration of Ydj1 in Y145X-Ydj1 droplets. Data shows mean ± SD for n = 3 independent experiments. (F) Lag time of Y145X aggregation via phase separation and with increasing Ydj1 concentrations. The lag times were estimated by fitting the ThT fluorescence kinetics to the sigmoidal function. AFM images with height profiles of the species formed in the saturation phase of aggregation kinetics of (G) Y145X only droplets and (H) in the presence of 20 µM Ydj1.

### Ydj1 initiates the phase separation of monomeric Y145X and inhibits its *de novo* aggregation by redirecting it toward a non-amyloidogenic pathway

In order to investigate the role of Ydj1 on monomeric Y145X, we first co-incubated Ydj1 (20 µM) and Y145X (50 µM) at physiological pH under the condition at which both Y145X and Ydj1 do not phase separate individually (Fig. S3A). Upon addition of Ydj1 to monomeric Y145X, the solution spontaneously turned turbid, indicating the presence of micron-sized droplets, which was further confirmed by performing two-color Airyscan imaging using sparsely AlexaFluor594-labeled Ydj1 and 1% AlexaFluor488-labeled single-Cys variant of Y145X (W99C) (Fig. S3B). It is known that under stress conditions, the pH becomes acidic, and the chaperone levels are elevated to maintain protein homeostasis and prevent misfolding and aggregation of the protein (63, 64). Therefore, we next set out to study the complex phase separation of Y145X and Ydj1 at acidic pH (pH 6.5), and as expected, they exhibited co-phase separation (Fig. S3B). Given that previous studies have reported a mildly acidic pH of ∼ 6.5 facilitates the conversion of monomeric Y145X into oligomeric and fibrillar aggregates (40, 65), we conducted our further studies at pH 6.5 to gain insights into how the presence of the Ydj1 modulates the aggregation behavior of the Y145X under these specific environmental conditions. We mixed Y145X and Ydj1 at different stoichiometries and observed their phase separation using turbidity measurements and two-color Airyscan imaging. The turbidity rose with increasing concentrations of Ydj1 at a fixed Y145X concentration, which is further corroborated by our concentration-dependent confocal imaging (Fig. S3C, 4A). Next, in order to study the material properties of these droplets, we performed FRAP experiments using 1 % of AlexaFluor488-labeled Y145X (W99C) that revealed rapid and near-complete recovery for Y145X, indicating the mobility and liquid-like nature of the droplets (Fig. 4B). Further, to investigate the diffusional properties of the droplets formed, we conducted single-droplet FCS measurements using AlexaFluor488-labeled Y145X (W99C). In its monomeric dispersed form, Y145X displayed a diffusion time of ⁓ 0.18 ms. However, this diffusion time increased dramatically by 350-fold to ⁓ 67 ms within the droplets, indicating a highly crowded and viscous environment within the condensed phase (Fig. 4C, D). Next, we aimed to discern the underlying molecular drivers governing this co-phase separation. Given that heterotypic condensation arises from electrostatic interaction between oppositely charged components, we postulated that the electrostatic forces could potentially promote their complex heterotypic phase separation. To verify the role of electrostatic interactions, we carried out the phase separation assays as a function of increasing ionic strength by performing two-color confocal imaging. Consistent with our hypothesis, the addition of increasing amounts of salt resulted in a complete dissolution of these droplets at 100 mM NaCl, highlighting the role of electrostatic interactions in the heterotypic phase separation of Y145X and Ydj1 (Fig. 4E). As mentioned previously, Hsp40 is known to interact with the polypeptides through hydrophobic interactions (60). Therefore, to study the effect of hydrophobic interactions in heterotypic phase separation, we created the M112Stop (M112X) variant of PrP (PrP 23-112), which is devoid of the hydrophobic region (HR) and carried out phase separation of M112X (50 µM) and Ydj1 (20 µM) with increasing salt concentration. If the hydrophobic effect is critical for complex phase separation, we would expect to observe the dissolution of the droplets at a salt concentration lower than that observed for Y145X-Ydj1. As anticipated, the co-phase separation was lower at 50 mM NaCl and completely abolished at 75 mM NaCl, as observed from two-color confocal imaging, highlighting the role of hydrophobic interactions in the phase separation of Y145X and Ydj1 (Fig. 4F). Subsequently, to investigate the effect of heterotypic interaction within the complex coacervates on the aggregation of Y145X, we performed the aggregation kinetics measurements of monomeric Y145X and Y145X-Ydj1 droplets formed using ThT under the stirring condition. Y145X alone exhibited a conventional nucleation-dependent polymerization with an increase in ThT fluorescence over time. In contrast, the ThT kinetics assay of Y145X-Ydj1droplets revealed a concentration-dependent inhibition or abrogation of amyloid aggregation of Y145X as indicated by the significantly lower final ThT fluorescence intensities (Fig. 4G, H). This was further supported by performing AFM imaging that revealed fibrillar aggregates with a height of about 4 nm for Y145X only, whereas conglomerate-like protein clusters were observed in the presence of Ydj1. Taken together, these findings suggest that the heterotypic interactions between Y145X and Ydj1 within the condensates effectively prevent the aggregation of Y145X into amyloid fibrils (Fig. 4I, J). The degree of inhibition appears to be dependent on the relative concentrations of the two components, with higher Ydj1 concentrations leading to more pronounced suppression of Y145X aggregation. Since nucleic acids are known to influence the protein phase behavior, we next set out to investigate the impact of Ydj1 on the complex phase separation involving Y145X and RNA.

**Fig. 4.**
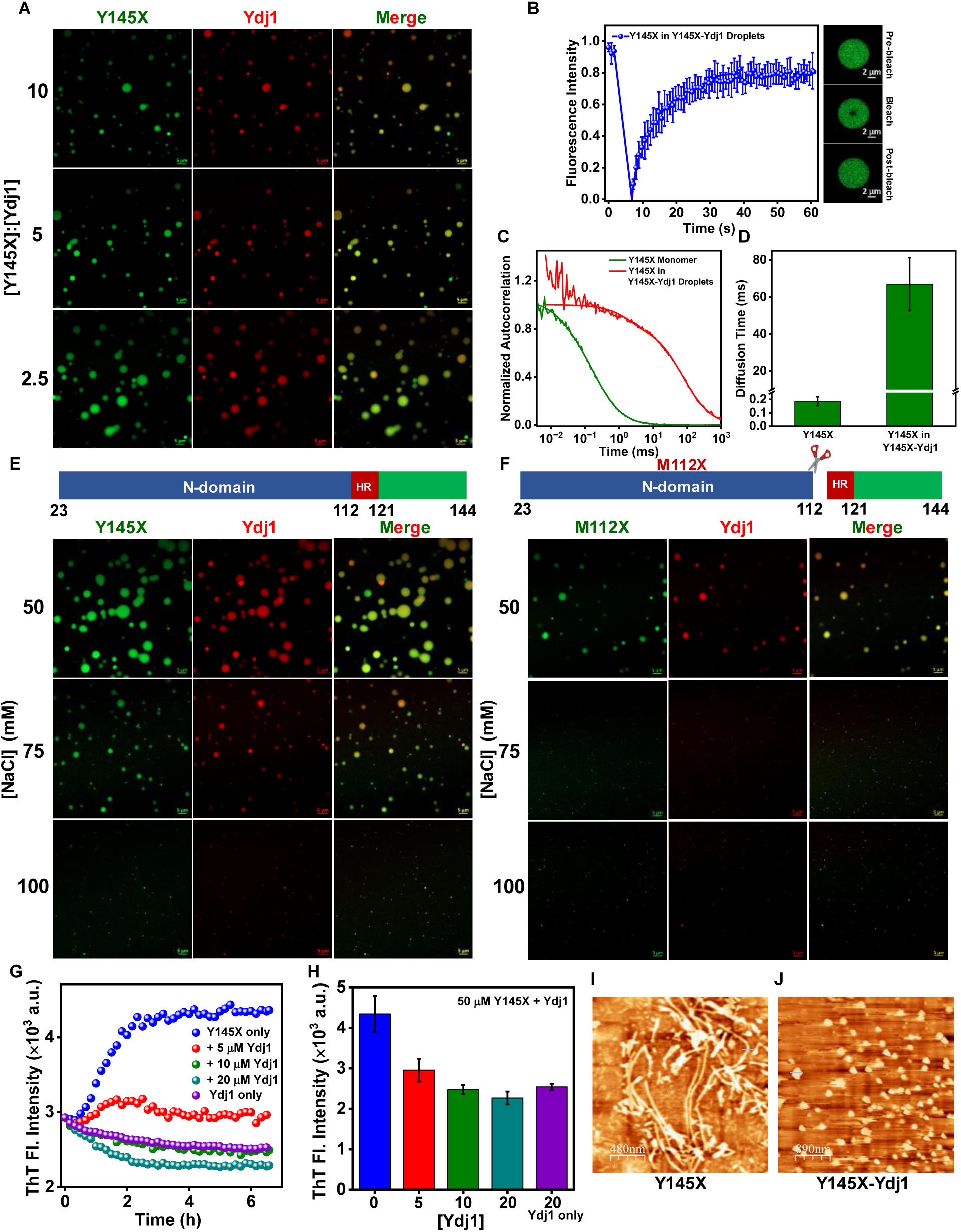
Ydj1 inhibits the aggregation of monomeric Y145X via complex coacervation. (A) Confocal microscopy images of AlexaFluor488-labeled Y145X (green) and AlexaFluor594-labeled Ydj1 (red) at different stoichiometries (scale bar 5 µm). (B) FRAP kinetics of Y145X in Y145X-Ydj1 complex coacervates using AlexaFluor488-labeled Y145X and fluorescence images of droplets obtained during FRAP measurements (scale bar 2 µm). Data represents mean ± SD of n = 5 independent experiments. (C) Normalized autocorrelation plots obtained from FCS measurements performed for monomeric Y145X and for Y145X in Y145X-Ydj1 droplets (unnormalized autocorrelation plots are shown in Fig. S3). (D) Diffusion time of Y145X in monomeric dispersed form, and in Y145X-Ydj1 droplets estimated from FCS measurements (Data represent mean ± SD for n = 5 independent samples). (E) Confocal images of 50 µM Y145X (AlexaFluor488-labeled, green) and 20 µM Ydj1 (AlexaFluor594-labeled, red) droplets with increasing salt concentrations (scale bar 5 µm). (F) Confocal images of 50 µM M112X (AlexaFluor488 NHS ester-labeled) and 20 µM Ydj1 (AlexaFluor594-labeled) droplets with increasing salt concentrations (scale bar 5 µm). (G) Representative ThT kinetics of *de novo* aggregation of Y145X-only and via phase separation mediated with increasing concentration of Ydj1 in Y145X-Ydj1 droplets. Data shows the mean for n ≥ 3 independent experiments. (H) Plot showing the final ThT intensity at the end of aggregation with increasing Ydj1 concentration. Data represents the mean ± SD for n ≥ 3 independent experiments. AFM images of the species formed in the saturation phase of aggregation kinetics of (I) Y145X only and (J) droplets formed in the presence of 20 µM Ydj1.

### Competitive binding within the Y145X-RNA-Ydj1 ternary system dictates the properties of multiphasic condensates

Biomolecular condensates are composed of diverse molecular species, including proteins, nucleic acids, and other components. The composition is often dynamic, allowing condensates to form, dissolve, and remodel in response to cellular signals and environmental changes. RNA, primarily due to its multivalent binding capacity, structural diversity, sequence specificity, flexibility, and regulatory roles, can act as a scaffold and can introduce multivalency in a multi-component system (37, 66). The N-terminal IDR of PrP has been shown to interact with RNA due to the presence of two highly conserved polybasic regions that contain clusters of lysine residues (35, 67–69). It has also been reported that the Y145X variant of PrP undergoes condensation in the presence of polyU RNA and results in the formation of less dynamic gel-like droplets (Fig. S5A) (33). Therefore, we next set out to see the effect of the Ydj1 on Y145X condensates formed in the presence of polyU RNA. We started by labeling the 3’ end of polyU RNA with AlexaFluor488-Hydrazide dye using periodate chemistry (Fig. S5B) (70). We then performed three color Airyscan fluorescence imaging with increasing polyU concentration (25-200 ng/µL), keeping Y145X (100 µM) and Ydj1 (10 µM) concentration constant and observed that the resulting droplets significantly varied in their internal organization and showed heterogeneous localization of RNA (Fig. 5A, S5C). These assemblies acquire immiscible multiphasic nested morphology, and the nested droplet structure persisted despite the variations in the RNA concentration (71). The formation of these multiphasic nested condensates was independent of the order in which the components were added to the reaction mixture. In this multiphasic system, Y145X interacting with RNA gets concentrated in the center while Y145X co-partitioning with Ydj1 was at the periphery, which was further supported by the Pearsons’ coefficient colocalization and Manders’ overlap analysis plots (Fig. 5B-D). This behavior can be ascribed to the competitive binding of RNA and Ydj1 with Y145X. Our FRAP experiments showed the slower and lesser recovery of Y145X at the center compared to the periphery, indicating slower diffusion of Y145X colocalizing with RNA and a highly dynamic nature at the periphery, suggesting competitive binding within the nested, multiphasic biomolecular condensates formed by Y145X, RNA, and Ydj1 (Fig. 5E, F). This sequestration of Ydj1 at the outer shell of the droplets can potentially regulate the phase behavior of the condensates formed. Since the condensates formed at low RNA concentrations are more rigid with a gel-like character, they can promote the formation of undesirable, insoluble aggregates over time. Therefore, we then aimed to investigate the crucial role of Ydj1 in modulating the maturation of Y145X condensates formed in the presence of RNA into solid-like aggregates. We monitored the maturation of RNA-induced droplets in the absence and presence of Ydj1 using ThT under agitation conditions. Our Y145X-RNA-Ydj1 ternary system did not show any measurable increase in the ThT intensity in contrast to RNA-induced droplets, indicating Ydj1-mediated abrogation of aggregate formation (Fig. 5G). Taken together, these findings indicate the importance of Ydj1 in preventing RNA-mediated aggregation of Y145X.

**Fig. 5.**
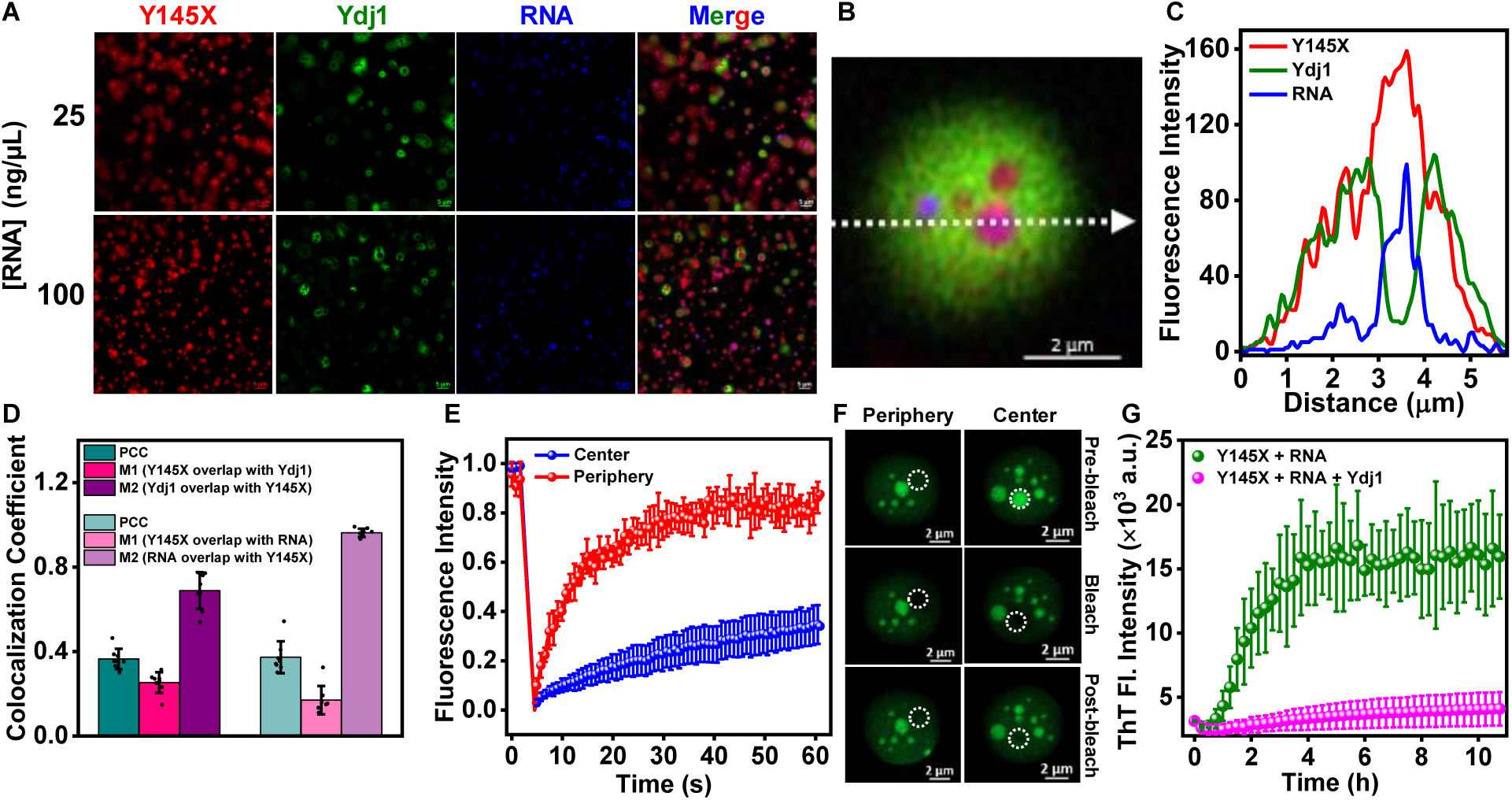
Ydj1 regulates Y145X-RNA condensates. (A) Two-color Airyscan confocal images of the Y145X-RNA-Ydj1 (AlexaFluor647-labeled Y145X, red; AlexaFluor594-labeled Ydj1, green; AlexaFluoro488-labeled RNA, blue) ternary system as a function of increasing RNA concentrations (Scale bar, 5 μm). Red, green, and blue colors are used for representation. (B) Representative image of the nested droplets formed by the Y145X-RNA-Ydj1 ternary system. (C) Representative fluorescence intensity profiles showing the competing interaction of Ydj1 and PolyU RNA with Y145X. (D) Pearson’s and Mander’s colocalization coefficient plots quantifying the fraction of Y145X co-partitioning with Ydj1 and vice-versa and Y145X co-partitioning with RNA and vice versa. (The data represent mean ± SD; n ≥ 15 high-resolution confocal images) (Y145 and Ydj1 concentration was 100 µM and 10 µM respectively, while RNA concentration was 25 ng/µl). (E) FRAP kinetics of Y145X colocalizing with RNA (center, blue) and Ydj1 (periphery, red) using AlexaFluor488-labeled Y145X single-Cys mutant at residue 31, (n ≥ 9) in the Y145X-RNA-Ydj1 ternary system. (F) Fluorescence images of droplets during FRAP measurements (area bleached is highlighted with the white dotted circle) (G) ThT kinetics of LLPS mediated aggregation of Y145X-RNA droplets (olive) and Y145X-RNA droplets in the presence of Ydj1 (pink).

## Discussion

Our study delves into the chaperone-mediated phase transition of a disease-associated amber-stop codon mutant of PrP (Fig. 6). We found that Ydj1 gets recruited into the phase-separated droplets of Y145X. This recruitment of Ydj1 into Y145X condensates resulted in liquid-like droplets, having a highly dynamic internal organization with the slower diffusion of Y145X in these droplets, indicating the formation of a dense viscoelastic network inside these heterotypic condensates. Our findings suggested that this partitioning of Ydj1 in Y145X droplets is driven by a complex interplay of hydrophobic and electrostatic interactions. Upon aging, we observed Ydj1-mediated inhibition of Y145X aggregation in a dose-dependent manner, kinetically trapping it into small non-amyloidogenic heteroprotein clusters by interacting with the aggregation-promoting region of Y145X. Taken together, our results showcase the crucial role of Hsp40 in regulating the phase transition of Y145X droplets and its assembly into fibrils, preventing the formation of insoluble amyloid-like aggregates. Furthermore, we demonstrated the effect of Ydj1 on the monomeric Y145X. Our findings suggest that Ydj1 potentiates the phase separation of Y145X via complex coacervation, leading to the formation of highly dynamic liquid-like droplets with a highly mobile internal organization at physiological and acidic pH. Our findings highlighted the role of Ydj1 in inhibiting the aggregation of monomeric Y145X via heterotypic coacervation by dynamically arresting them into nanoscopic hetero-clusters of Y145 and Ydj1 and preventing it from accessing the aggregation-prone state. This phase separation-mediated complex coacervation of Y145X with Ydj1 can potentially provide a mechanistic underpinning of chaperone-mediated regulation. Our findings emphasize the crucial role of Hsp40 in maintaining the metastable state of membrane-less organelles and preventing aberrant protein aggregation associated with neurodegenerative disorders. Our results further revealed the crucial role of Ydj1 in modulating the material properties of the phase-separated condensates formed in the presence of cofactors such as RNA by forming a ternary system of nested droplets with a distinct spatial organization. The sequestration of Ydj1 at the outer shell of the nested droplet can potentially regulate the phase transition of gel-like condensates formed in the presence of RNA by preventing the uncontrolled solidification and aggregation, thereby acting as a stabilizer, even in the presence of a cofactor like RNA.

**Fig. 6.**
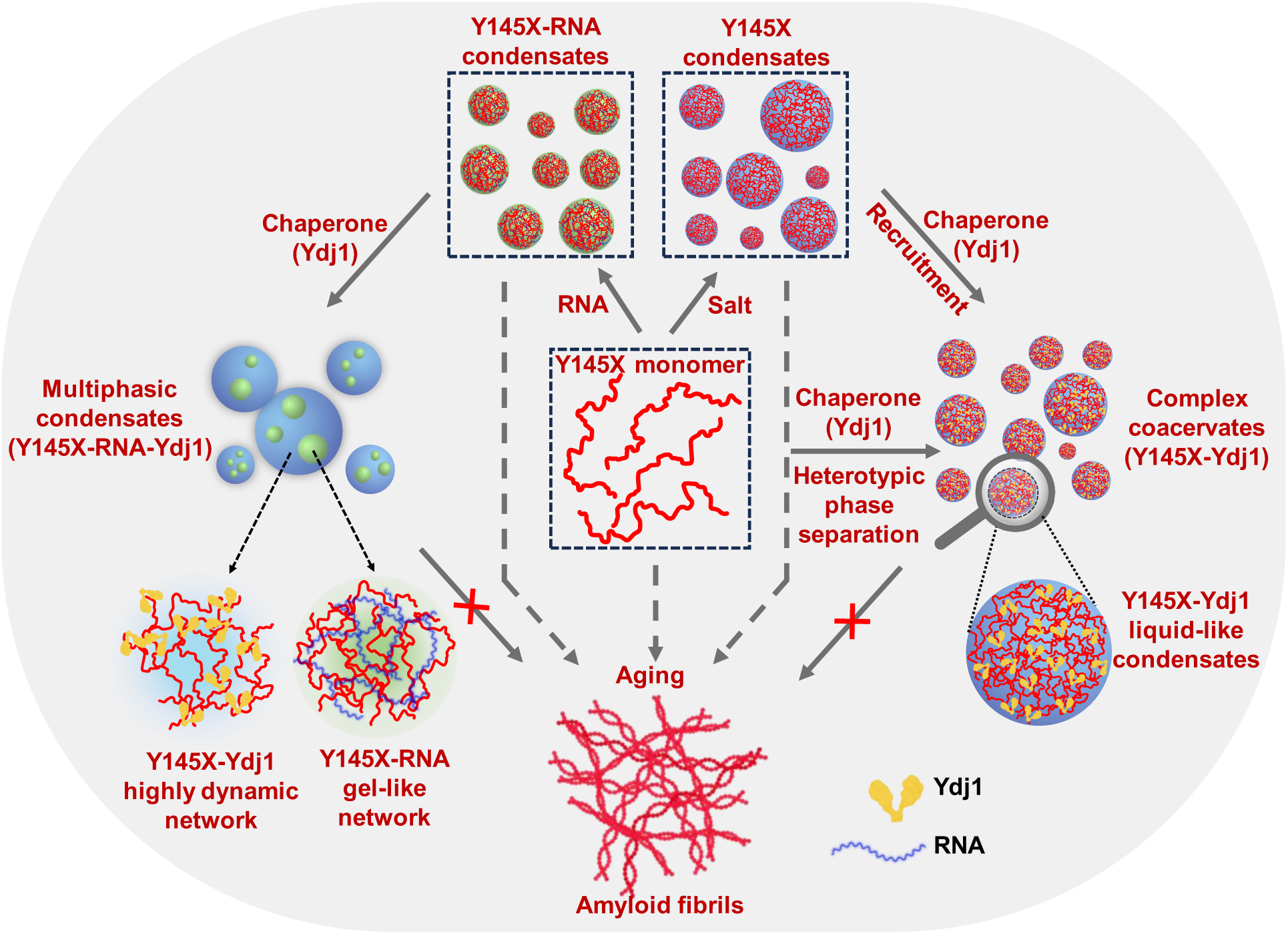
Schematic illustration of the role of Ydj1 in the regulation of phase transition of Y145X. Monomeric Y145X and droplets of Y145X formed in the presence of cofactors such as salt and RNA undergo a liquid-to-solid transition into amyloid-like aggregates associated with neurodegeneration. Ydj1 modulates the phase behavior of Y145X by binding to the charge clusters and hydrophobic region of Y145X, thereby inhibiting its aggregation.

In summary, our findings present the important role of Ydj1 in stabilizing the phase-separated condensates by regulating the phase separation and aggregation of disease-associated Y145X that readily undergoes phase separation-mediated aggregation (Fig.6). Cellular PQC is an intricate process that requires the synchronized effort of multiple chaperones operating in a feedback loop to eliminate toxic protein species and pathological aggregates. Under normal conditions, the misfolded forms of PrP are degraded in cells by the endoplasmic reticulum (ER)-associated degradation (ERAD)-proteasome pathway. However, upon proteasomal inhibition, the proteins fated to undergo degradation accumulate in the cytoplasm and assemble into toxic aggregates potentially leading to pathogenesis (41, 72). Although previous studies have highlighted the role of ER chaperones and other Hsps (Hsp60, Hsp70, and various small pharmacological chaperone molecules) in modulating PrP-induced toxicity and prion propagation either by stabilizing the native state or suppressing interactions driving aggregation, the role of Hsp40 as an independent chaperone in regulating the aggregation of Y145X via phase separation is important (73). Our work underscores the crucial role of Hsp40 chaperones as independent regulators of protein phase separation and aggregation within the cellular PQC network. This property of phase separation of Hsp40 with other misfolded proteins in membraneless organelles may be crucial for facilitating chaperone-mediated protein homeostasis within these biological condensates. Previous studies have also highlighted the role of small Hsps and Hsp40 on the phase separation and aggregation of FUS, suggesting its potential role in other neurodegenerative diseases (49, 58). Our findings provide mechanistic underpinnings of how molecular chaperones can act as independent regulators of the protein phase behavior with broader implications for the maintenance of cellular proteostasis and the prevention of protein aggregation associated with fatal neurodegenerative diseases.

## Material and Methods

Detailed materials and methods are provided in Supporting Information. The human Y145X construct was generated using full-length human PrP (23–231) plasmid. All the mutations, including single cysteine variants of Y145X and A→V Y145X mutant, were created using a QuikChange site-directed mutagenesis kit (Stratagene), as mentioned previously. Cleavable N-terminal His-tagged Y145X was purified using a Ni-NTA column, followed by His-tag removal with thrombin. Cleavable N-terminal His-tagged Ydj1 was purified using a Co-NTA column, and the His-tag was removed by TEV protease. Phase separation assays were conducted using turbidity measurements and confocal microscopy. FRAP experiments were performed using 0.2 % AlexaFluor488-C5-maleimide–labeled protein on a Zeiss LSM 980 equipped with an Elyra 7 super-resolution microscope. FCS experiments and single-droplet anisotropy measurements were performed using AlexaFluor488-C5-maleimide-labeled and F5M-labeled Y145X protein, respectively, on the MT200 time-resolved fluorescence confocal microscope (PicoQuant). Aggregation kinetics were performed using NUNC 96-well plates on a POLARstar Omega Plate Reader Spectrophotometer (BMG Labtech). Vibrational Raman spectroscopic studies were conducted on an inVia laser Raman microscope (Renishaw) using a 100× objective lens (Nikon) and a 785-nm near-infrared laser. AFM images were acquired on an Innova atomic force microscope (Bruker) operating in tapping mode.

## Acknowledgments

We thank IISER Mohali, Department of Science and Technology, Govt. of India (FIST grant # SR/FST/LS-II/2017/97 to the Department of Biological Sciences, IISER Mohali), Science and Engineering Research Board (SUPRA SPR/2020/000333, CRG/2021/002314, and J.C. Bose Fellowship JCB/2023/000016 to S.M.), Ministry of Education, Govt. of India (Centre of Excellence grant to S.M.), and Indo-French Centre for the Promotion of Advanced Research (IFC/A/6903-3/2023/680 to S.M.), Prof. Witold Surewicz (Case Western Reserve University) and Prof. Deepak Sharma (Institute of Microbial Technology, Chandigarh) for the kind gift of the DNA plasmids for full-length prion protein and Ydj1, respectively, Prof. N. Periasamy (Retd. TIFR Mumbai) for providing us with the fluorescence decay analysis program, and the members of the Mukhopadhyay lab for critically reading this manuscript.

## Authors Contributions

L.A. and S.M. conceived the project. L.A., D.B., S.S., S.K.R., and A.S. performed the experiments and analyses. L.A. and D.B. prepared the figures, and L.A. wrote the first draft. S.M. supervised the work, edited the manuscript, obtained funding, and provided the overall direction. All authors discussed the results and commented on the manuscript.

