## Supplementary Information for "Chaperone-mediated heterotypic phase separation prevents the amyloid formation of the pathological Y145Stop variant of the prion protein"

[§]Contributed equally

*Corresponding author:

Samrat Mukhopadhyay

### Materials and Methods

#### Materials

Sodium phosphate monobasic dihydrate, sodium phosphate dibasic dihydrate, tris base, 2-mercaptoethanol, sodium hydroxide, sodium acetate, sodium periodate, sodium chloride, magnesium chloride, potassium chloride, imidazole, Thioflavin T, 1,4-dithiothreitol (DTT), PolyU sodium salt, phenylmethylsulfonyl fluoride (PMSF), TCEP (tris(2-carboxyethyl)phosphine), ethylenediaminetetraacetic acid (EDTA), nuclease-free water (NFW), were of highest purity grade, procured from Sigma (St. Louis, MO, USA). Guanidinium hydrochloride and urea were purchased from SRL and Himedia, respectively. Ampicillin, chloramphenicol, and isopropyl-β-thiogalactopyranoside (IPTG) were obtained from Gold Biocom (USA). NUNC 96-well plates were procured from ThermoFisher Scientific (Waltham, Massachusetts, U.S.A). All the fluorescent probes used in the experiments, namely, fluorescein-5-maleimide (F5M), AlexaFluor488-C5-maleimide, AlexaFluor488-hydrazide, and AlexaFluor594-C5-maleimide, were purchased from Molecular Probes, Invitrogen. Ni-NTA resin was procured from Qiagen, and Co-NTA resin was obtained from G-Biosciences. NAP-10 and NAP-5 columns were procured from GE Healthcare Life Sciences (USA). Amicon membrane filters for concentrating proteins were purchased from Merck Millipore.

#### Methods

##### Bioinformatic analysis

FuzDrop was used to determine the phase separation propensity of Y145X and Ydj1 (http://protdyn-fuzpred.org/) (1). The Classification of Intrinsically Disordered Ensemble Regions (CIDER) tool was used to determine the net charge distribution throughout the protein polypeptide chains and their linear hydropathy profile (http://pappulab.wustl.edu/CIDER/analysis) (2).

##### Construct details and site-directed mutagenesis

The Y145X construct was made using human PrP (23-231) plasmid as a template, which was a kind gift from Prof. W. K. Surewicz (Case Western Reserve University, USA). The single-cysteine mutants and A→V Y145X were created using the QuickChange kit (Stratagene), as described previously (3, 4). The M112X variant of human PrP was created by site-directed mutagenesis (KOD Master Mix) using Y145X plasmid as a template (Table S1). All the mutations were verified by Sanger sequencing. The plasmid coding Ydj1 was a kind gift from Prof. Deepak Sharma (Institute of Microbial Technology, Chandigarh, India).

##### Recombinant protein expression and purification

Proteins were expressed and purified following previously published protocols, with slight modifications (5, 6). The wild-type, all the other mutants of Y145X, and M112X were overexpressed in BL21(DE3)pLysS cells and purified using Nickel-NTA affinity chromatography. Briefly, bacterial cultures of Y145X, all the other mutants of Y145X and M112X were induced with 1 mM IPTG at O.D.600 = 0.6 for 8 hours at 30 °C. Cell pellets were harvested by centrifugation at 4 °C, 3220 x g for 35 minutes. Purification of cleavable His-tagged constructs of Y145X and M112X was performed using Ni-NTA chromatography as previously described (5). The cysteine mutants of Y145X were purified by resuspending the cell pellets in lysis buffer (50 mM Tris, 100 mM NaCl, 1 mM EDTA, 0.1 % Triton X-100), followed by sonication using a probe sonicator (5% Amplitude, 15 seconds ON, 10 seconds OFF for 25 minutes). The inclusion bodies were isolated by high-speed centrifugation, washed, and resuspended in the denaturation buffer. The dissolved pellet was subjected to incubation for 2 h, followed by high-speed centrifugation. The supernatant was loaded onto a pre-equilibrated Ni-NTA column, and the bound protein was then eluted with a buffer containing 500 mM imidazole (8 M urea, 10 mM tris, 100 mM sodium phosphate, pH 7.5). Following the purification, the purified N-terminal His-tagged protein was dialyzed overnight at room temperature against a native dialysis buffer (20 mM sodium phosphate, 50 mM NaCl, pH 6.5). The N-terminal 6X-His tag was cleaved using 0.2 U/mL thrombin protease at 37 °C for 5 h. After the cleavage reaction, the thrombin protease was inactivated using 0.2 mM PMSF. Further, to separate cleaved and uncleaved fractions, the protein was loaded onto a Ni-NTA column and eluted with 20 mM imidazole buffer (10 mM Tris, 100 mM sodium phosphate, 8 M urea, pH 8.0). The pure protein was further concentrated using 3 kDa MWCO Amicon membrane filters and buffer exchanged into phosphate buffer (20 mM phosphate buffer, pH 7.5 or pH 6.5) using the NAP-10 column. The purity of the protein was determined using SDS-PAGE gel analysis, and pure protein concentration was estimated using $\varepsilon_{280}$ = 43,670 $M^{-1}cm^{-1}$ for Y145X and $\varepsilon_{280}$ = 42390 $M^{-1}cm^{-1}$ for M112X. All the experiments were performed using freshly purified proteins to avoid freeze-thaw cycles.

Recombinant N-terminal 6X-His-Ydj1 cloned in the pPROEX-HTb vector was transformed into BL21(DE3) Rosetta cells, overexpressed, and purified using cobalt-NTA affinity chromatography. Bacterial cultures were grown at 37 °C, 220 rpm to an O.D.600 = 0.6. The overexpression was induced using 0.3 mM IPTG at 16 °C for 16 h. Then, cells were harvested by centrifugation at 4 °C, 3220 x g for 35 min. The harvested cell pellets were immediately resuspended in ice-cold lysis buffer (25 mM HEPES, 500 mM NaCl, 20 mM KCl, 20 mM $MgCl_2$, pH 7.4) and were incubated with lysozyme (2 mg/ml) at 4 °C. Then sonication was performed, followed by high-speed centrifugation at 4 °C, 15,557 x g, to get rid of the cell debris. The supernatant was loaded onto a pre-equilibrated Co-NTA column, and further gradient washes of imidazole were given. Finally, the protein was eluted in elution buffer (25 mM HEPES, 500 mM

NaCl, 20 mM KCl, 20 mM $MgCl_2$, 300 mM imidazole, 1 mM DTT, pH 7.4). The N-terminal 6X-His tag was cleaved overnight at 4 °C in dialysis buffer (25 mM HEPES, 500 mM NaCl, 20 mM KCl, 20 mM $MgCl_2$, 1 mM DTT, pH 7.4) using recombinantly purified Tobacco Etch Virus (TEV) protease. Further, cleaved Ydj1 was passed through Co-NTA to remove His-tag and un-cleaved proteins. The eluted pure protein was concentrated using a 10 kDa MWCO Amicon membrane filter, and concentration was estimated by measuring absorbance at 280 nm ($\varepsilon$ = 23,475 $M^{-1}$ $cm^{-1}$). The purity of the protein was confirmed by running an SDS-PAGE. Small aliquots were made, flash-frozen, and stored at – 80 °C.

**Phase separation assays**

Phase separation of Y145X (100 μM) was immediately induced by adding NaCl (350 mM) to the reaction mixture in 20 mM sodium phosphate buffer, pH 7.5. For all the co-partitioning experiments, varying concentrations of Ydj1 were added to these preformed droplets of Y145X. For the rest co-phase separation experiments, 50 μM Y145X was mixed with varying concentrations of Ydj1 in 20 mM sodium phosphate, 50 mM NaCl buffer, pH 6.5. The immediate rise in turbidity indicated the complex coacervation of Y145X and Ydj1. The turbidity measurements were performed by recording the absorbance at 350 nm using a Multiskan Go (Thermo scientific) plate reader using a 96-well NUNC optical bottom plate. The reaction volumes were kept constant at 120 μL. The mean and standard error were obtained from at least three independent sets of measurements on the same day.

**Fluorescence labeling**

All the cysteine mutants of Y145X were labeled using thiol-reactive maleimide chemistry under denaturing buffer conditions (10 mM Tris, 100 mM sodium phosphate, 8 M urea, pH 7.4). For labeling with F5M dye, 20-fold molar excess dye was added, and for AlexaFluor dyes, 2-fold molar excess dye was added. First, the purified protein was incubated with 0.3 mM tris(2-carboxyethyl)phosphine (TCEP) for 30 mins, and then the dye was added to the reaction mixture. The reaction was incubated for 2-3 h in the dark at room temperature under stirring conditions. Excess-free dye was removed using a NAP-10 column. Ydj1 was labeled using AlexaFluor dyes in 1:1 (Protein: Dye) molar ratio under native buffer conditions (25 mM HEPES, 500 mM NaCl, 20 mM $MgCl_2$, 20 mM KCl, 0.3 mM TCEP, pH 7.4). Excess unreacted dye was removed using a 10 kDa MWCO Amicon filter. The labeled protein concentration was estimated using $\varepsilon_{495}$ = 68,000

$M^{-1}cm^{-1}$ for F5M, $\varepsilon_{495}$ = 72,000 $M^{-1}cm^{-1}$ for AlexaFluor488-C5-maleimide, $\varepsilon_{590}$ = 92,000 $M^{-1}cm^{-1}$ for AlexaFluor594-C5-maleimide, and $\varepsilon_{647}$ = 2,39,000 $M^{-1}cm^{-1}$ for AlexaFluor647-C2-maleimide. For RNA labeling, the 3′ end of polyU was labeled with AlexaFluor488 Hydrazide, as described previously (7). Firstly, the 2′-3′ diols of RNA were oxidized into aldehyde by sodium periodate, followed by precipitation. Resuspended RNA was incubated with 20-fold molar excess AlexaFluor488 hydrazide dye in 0.1 M sodium acetate, pH 5.2 buffer conditions. The labeling reaction was incubated at 25 °C under stirring conditions, followed by incubating it overnight at 4 °C. The excess unreacted free dye was removed using a NAP-5 column. Labeled RNA was precipitated and eventually resuspended in DEPC-treated water. The labeled RNA concentration was estimated using a TECAN Infinite Pro 200 microplate plate reader.

**Estimation of saturation concentration ($C_{sat}$)**

The Y145X only and Y145X-Ydj1 droplet reactions doped with 1 % AlexaFluor488 labeled Y145X were set and incubated at room temperature for 10 mins. The reactions were then subjected to high-speed centrifugation at 25 °C, 18,000 x g for 30 min. The supernatant was removed carefully without disturbing the pellet. The dilute phase protein concentration was estimated by monitoring absorbance at 495 nm and a factor of 100 was incorporated for $C_{sat}$ calculations.

**Confocal microscopy**

Fluorescence imaging experiments were performed on a ZEISS LSM 980 Elyra 7 super-resolution microscope, having a high-resolution monochrome cooled AxioCamMRm Rev. 3 FireWire (D) camera, using a 63x oil-immersion objective (Numerical aperture 1.4). The droplet reaction was incubated at room temperature for 5 min, and ~ 10 µL of the sample was placed on a glass coverslip and imaged using Airyscan 2 detector (32 channels GaAsP). Reactions doped with 1% labeled protein were imaged using a 488 nm laser diode (11.9 mW), a 590 nm excitation source, and a 632 nm excitation source for AlexaFluor488, AlexaFluor594, and AlexaFluor647 labeled proteins respectively. Images were acquired at 1840 x 1840 pixels and 16-bit depth resolution. All the fluorescence images were processed and analyzed using the instrument software Zen Blue 3.2.

**Fluorescence recovery after photobleaching (FRAP) measurements**

FRAP measurements were performed on ZEISS LSM 980 Elyra 7 super-resolution microscope using a 63x oil-immersion objective (Numerical Aperture 1.4) and a monochrome cooled high-resolution AxioCamMRm Rev. 3 FireWire (D) camera. AlexaFluor488-C5-maleimide labeled protein (1%) was used for FRAP experiments. For image acquisitions, processing, and acquiring FRAP traces, ZEN (2011) software was used. For all the experiments, a region of interest (ROI) with a radius of 1 µm for droplets at pH 7.5 and 0.5 µm for droplets at pH 6.5, and in the presence of RNA was bleached using a 488 nm laser diode. The recovery was recorded using the Zen Blue 3.2 (ZEISS) software. The recovery traces were background corrected, normalized, and plotted using Origin (2021b). The plots were fitted using a single-exponential equation to determine the half-life of recovery ($t_{1/2}$).

**Single-droplet steady-state and time-resolved fluorescence anisotropy measurements**

Fluorescence anisotropy measurements were recorded using the PicoQuant MicroTime 200 (MT200) time-resolved confocal microscope. Measurements were performed of monomeric Y145X, Y145X only droplets, and Y145X-Ydj1 droplets spiked with 0.1 % F5M labeled single cysteine variants of Y145X at residue positions 31, 99, 120. Fresh droplet reactions (50 µL) were set and dropped over a glass coverslip placed directly on a Super Apochromat 60x objective (water immersion) with 1.2 NA (Olympus). A 485 nm laser was used to excite the samples, and emitted fluorescence was selectively collected using a 520/35 bandpass filter and passed through a 50 µm pinhole to get rid of out-of-focus light. The emitted photons were split into two SPAD (Single-Photon Avalanche Diode) detectors using a polarizing beam splitter. Anisotropy imaging was performed for steady-state information, while point time traces were acquired for time-resolved measurements. Correction factors were estimated using a free dye solution, utilizing which the steady-state anisotropy values were estimated using the commercially available SymphoTime64 software v2.7. The steady-state anisotropy is given by the following equation:

$$r_{ss} = \frac{I_{\parallel} - I_{\perp}}{[1-3L2]I_{\parallel} + [2-3L1]I_{\perp}} \quad (1)$$

Where $I_{\parallel}$ and $I_{\perp}$ are the background corrected parallel and perpendicular fluorescence intensities and L1 and L2 are the correction factors for the used objective lens.

For time-resolved decay analysis, the obtained decay profiles were further fitted globally using the following relations:

$$I_{\parallel}(t) = 1\,/\,3I(t)[1 + 2r(t)] \quad (2)$$

$$I_{\perp}(t) = 1\,/\,3I(t)[1 - r(t)] \tag{3}$$

where *I* represent the time-dependent fluorescence intensity recorded at the magic angle (54.7°). The time-resolved fluorescence anisotropy decay profiles were fitted using a biexponential decay function and can be expressed as follows:

$$r(t) = r_0\left[\beta_1 e^{\left(\frac{-t}{\phi_1}\right)} + \beta_2 e^{\left(\frac{-t}{\phi_2}\right)}\right] \tag{4}$$

Where $\phi_1$ and $\phi_2$ represent the fast and slow rotational correlation times associated with the local and global dynamics of the protein chain, respectively. β1 and $\beta_2$ are the amplitudes corresponding to $\phi_1$ and $\phi_2$ rotational correlation times. $r_0$ stands for the time-zero fundamental anisotropy of the attached fluorophore. The goodness of fit was estimated using the reduced $\chi^2$ values (8).

**Fluorescence correlation spectroscopy (FCS)**

FCS experiments were performed using the instrument MicroTime 200 for the monomeric Y145X and Y145X in the droplet phase in the absence and presence of Ydj1. A stock of 1 nM AlexaFluor 488 free dye solution was used to estimate the confocal volume (Veff) and the corresponding structural parameter (κ) for our setup. Using the estimated parameters, Veff > 1fL and κ = 4.7, as calibration values, the acquired data were curve-fitted. Measurements were taken by focusing inside the solution (50 µm) in the case of dispersed monomers, whereas individual droplets were focused in the case of phase-separated reactions. The data were acquired and analyzed, and correlation curves were fitted with the triplet-state model using in-built SymphoTime64 software v2.7. The diffusion time of Y145X within the monomeric and condensed phases was obtained.

**Amyloid aggregation kinetics**

The aggregation kinetics via phase separation and classical nucleation-dependent pathway were monitored in the absence and presence of varying Ydj1 concentrations, using NUNC 96-well optical bottom plate on POLARstar Omega Plate Reader Spectrophotometer (BMG LABTECH, Germany) at room temperature. The reaction volume in each well was kept constant at 150 µL with a glass bead of 3 mm diameter. Thioflavin T (ThT) was used at a final concentration of 20 µM in the aggregation reactions. The kinetics were monitored under stirring conditions for 60 seconds before each cycle at 100 rpm. The acquired kinetics data was plotted using Origin (2021b).

**Raman spectroscopy**

All the Raman spectra were recorded on an inVia laser Raman microscope (Renishaw, UK) at 25 °C. The condensed phase of the droplet reactions was used for acquiring the spectra. After incubating the droplet reactions, it was subjected to high-speed centrifugation. The supernatant was removed carefully without disturbing the dense phase pellet, and the dense phase was resuspended in 6 µL sodium phosphate buffer (20 mM sodium phosphate, pH 7.5) and deposited on a glass slide covered with aluminum foil. The half-dried samples were focused using a 100x objective lens (Nikon, Japan) and excited using a 785 nm NIR laser, with an exposure time of 10 s and 100 % (500 mW) laser power. The Rayleigh scattering was filtered using an edge filter of 785 nm. The collected Raman scattering was dispersed using a 1200 lines/mm diffraction grating and detected using an air-cooled CCD detector (9). All the data were acquired for 10 accumulations using the in-built software Wire 3.4. Acquired spectra were baseline corrected and smoothened using Wire 3.4 and plotted using Origin (2021b).

**Atomic force microscopy (AFM) imaging**

AFM images were obtained using an Innova atomic force microscope (Bruker) operating in tapping mode. The sample was prepared by drop casting 10 µL reaction mixture onto freshly cleaved, Milli-Q water-washed muscovite mica (Grade V-4 mica from SPI, PA) followed by incubation at room temperature for 5 min and two 150 µL washes of Mili-Q water. The samples were further dried using a gentle stream of nitrogen gas. The images were acquired using NanoDrive (v8.03) software and processed using WSxM 5.0D 8.1 software. The height profiles were determined from WSxM and were plotted in the Origin software (2021b) (10).

**Table S1**. Primers used for creating M112X truncation.

| | |
|---|---|
| M112X Forward | ACCAACATGAAGCACTAGGCTGGTGCTGCAGCA |
| M112X Reverse | TGCTGCAGCACCAGCCTAGTGCTTCATGTTGGT |

**Table S2**. Recovered parameters from time-resolved fluorescence anisotropy decay analyses.

| **Y145X** | | **$\phi_1$ (ns)**<br>**($\beta_1$)** | **$\phi_2$ (ns)**<br>**($\beta_2$)** |
|---|---|---|---|
| **Residue 31** | Monomer | 1.12 ± 0.07<br>0.61 ± 0.03 | 4.45 ± 0.63<br>0.39 ± 0.03 |
| | Y145X Droplets | 1.22 ± 0.17<br>0.20 ± 0.02 | 53.59 ± 2.62<br>0.80 ± 0.02 |
| | Y145X-Ydj1 Droplets | 1.78 ± 0.11<br>0.23 ± 0.02 | 77.47 ± 5.40<br>0.77 ± 0.02 |
| **Residue 99** | Monomer | 0.95 ± 0.09<br>0.56 ± 0.03 | 4.38 ± 0.11<br>0.44 ± 0.03 |
| | Y145X Droplets | 1.42 ± 0.05<br>0.22 ± 0.02 | 65.30 ± 5.89<br>0.78 ± 0.02 |
| | Y145X-Ydj1 Droplets | 1.38 ± 0.12<br>0.22 ± 0.02 | 72.18 ± 7.56<br>0.78 ± 0.023 |
| **Residue 120** | Monomer | 1.01 ± 0.08<br>0.75 ± 0.04 | 4.57 ± 0.60<br>0.25 ± 0.04 |
| | Y145X Droplets | 1.28 ± 0.06<br>0.20 ± 0.01 | 78.81 ± 6.41<br>0.80 ± 0.01 |
| | Y145X-Ydj1 Droplets | 1.36 ± 0.10<br>0.21 ± 0.02 | 74.80 ± 9.83<br>0.79 ± 0.02 |

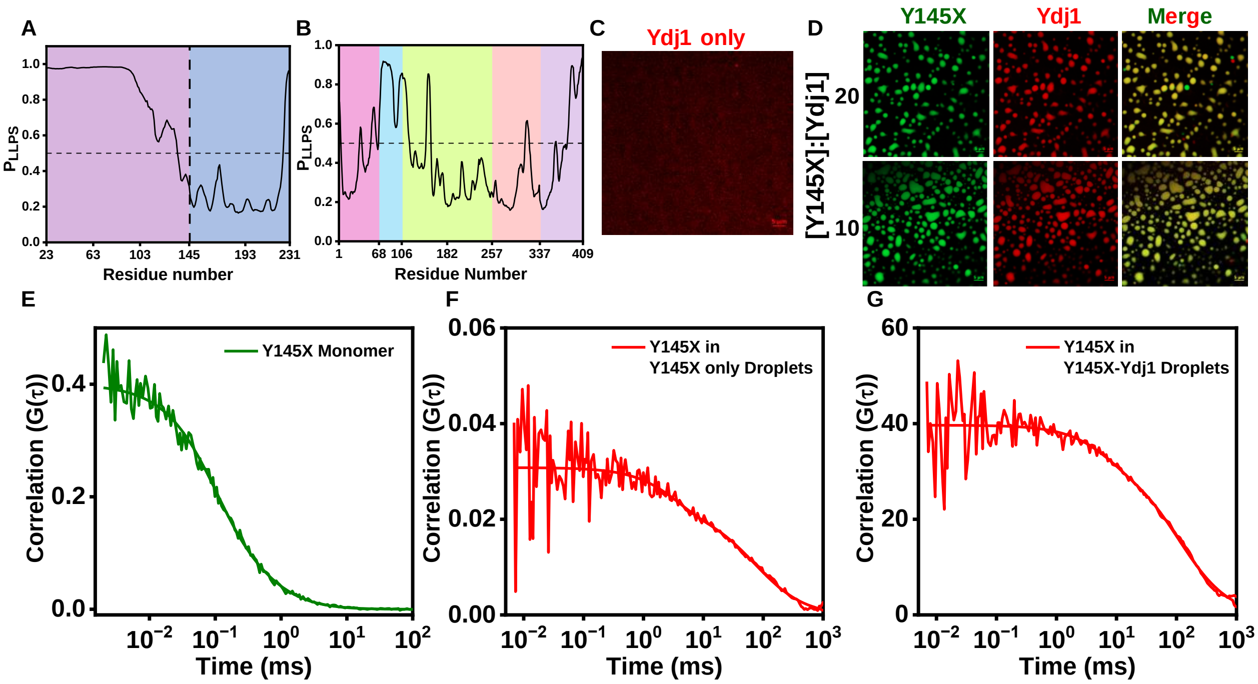


**Fig. S1.** Phase separation propensity of (A) Y145X and (B) Ydj1 using FuzDrop. (C) Confocal fluorescence image of 10 µM Ydj1 (AlexaFluor594-labeled, 20 mM sodium phosphate, 350 mM NaCl, pH 7.5) (scale bar 5 µm). (D) Two-color confocal microscopy images of heterotypic condensates of Ydj1 (AlexaFluor594-labeled, red) and Y145X (AlexaFluor488-labeled, green) at different stoichiometries as indicated, with Y145X concentration constant (100 µM) (scale bar 5 µm). Unnormalized FCS autocorrelation plots with fits of (E) Y145X monomer, (F) Y145X in Y145X only droplets, (G) Y145X in Y145X-Ydj1 droplets.

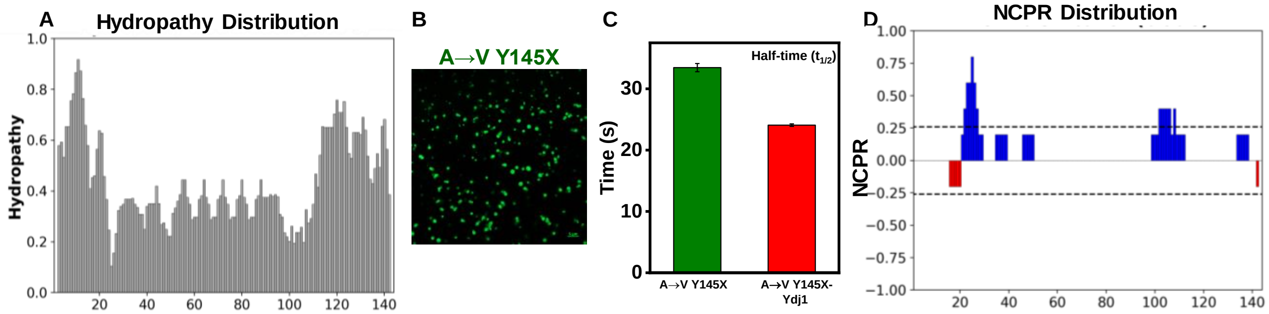


**Fig. S2.** (A) Linear hydropathy plot of Y145X (plotted using sequence from 1-144 residues) using CIDER. (B) Confocal image of droplets formed by A→V Y145X variant (100 µM protein, 20 mM sodium phosphate, 350 mM NaCl, pH 7.5). (C) Half-time ($t_{1/2}$) plot of A→V Y145X in A→V Y145X only droplets and in the presence of Ydj1. (D) Charge distribution profile (NCPR- Net charge per residue) of Y145X (plotted using the sequence from 1-144 residues). Red and blue colors indicate negatively charged and positively charged residues, respectively.

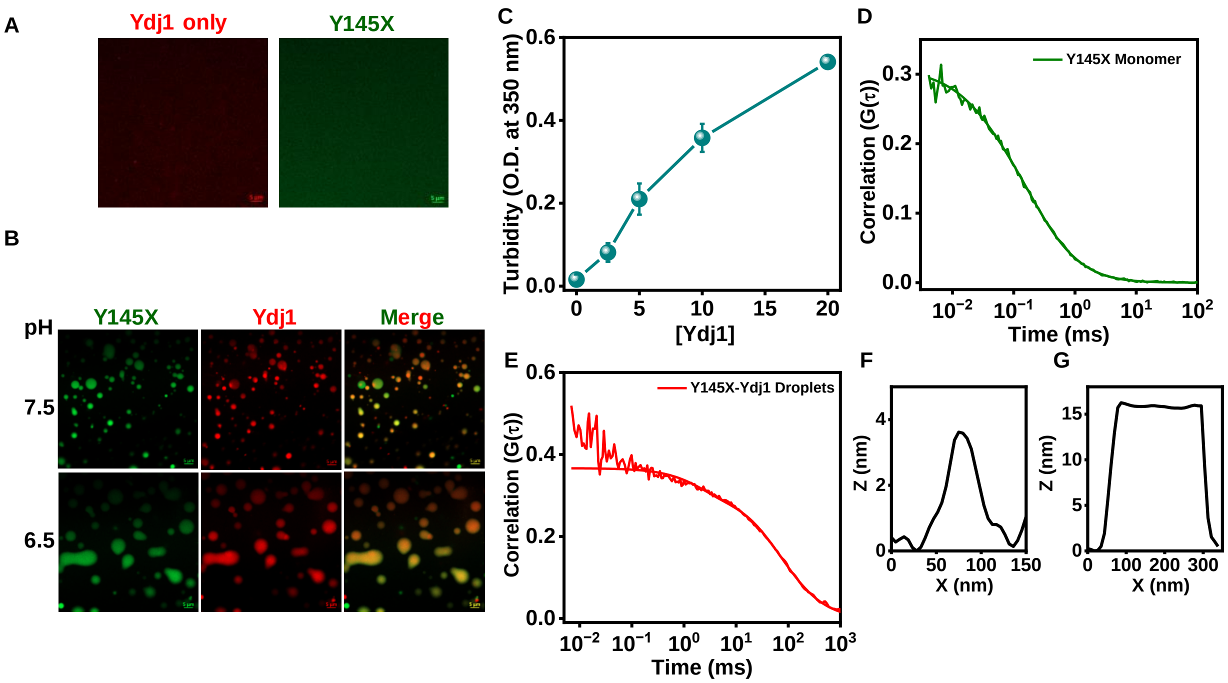


**Fig. S3.** (A) Confocal images of the dispersed phase of 20 μM Ydj1 (AlexaFluor594-labeled) and 50 μM Y145X (AlexaFluor488-labeled) (20 mM sodium phosphate, 50 mM NaCl, pH 6.5, scale bar 5 μm). (B) Two-color Airyscan images of Y145X-Ydj1 droplets formed at two different pH (Ydj1 – red, Y145X – green, scale bar 5 μm). (C) Concentration-dependent turbidity (at O.D. 350 nm) of Y145X-Ydj1 reaction mixtures. The data represent mean ± SD for n = 3 independent single-day measurements. Unnormalized FCS autocorrelation plots with fits of (D) Y145X monomer and (E) Y145X in Y145X-Ydj1 droplets. Height profiles obtained from AFM images of the species formed in the saturation phase of aggregation kinetics of (F) Y145X only and (G) droplets formed in the presence of 20 μM Ydj1.

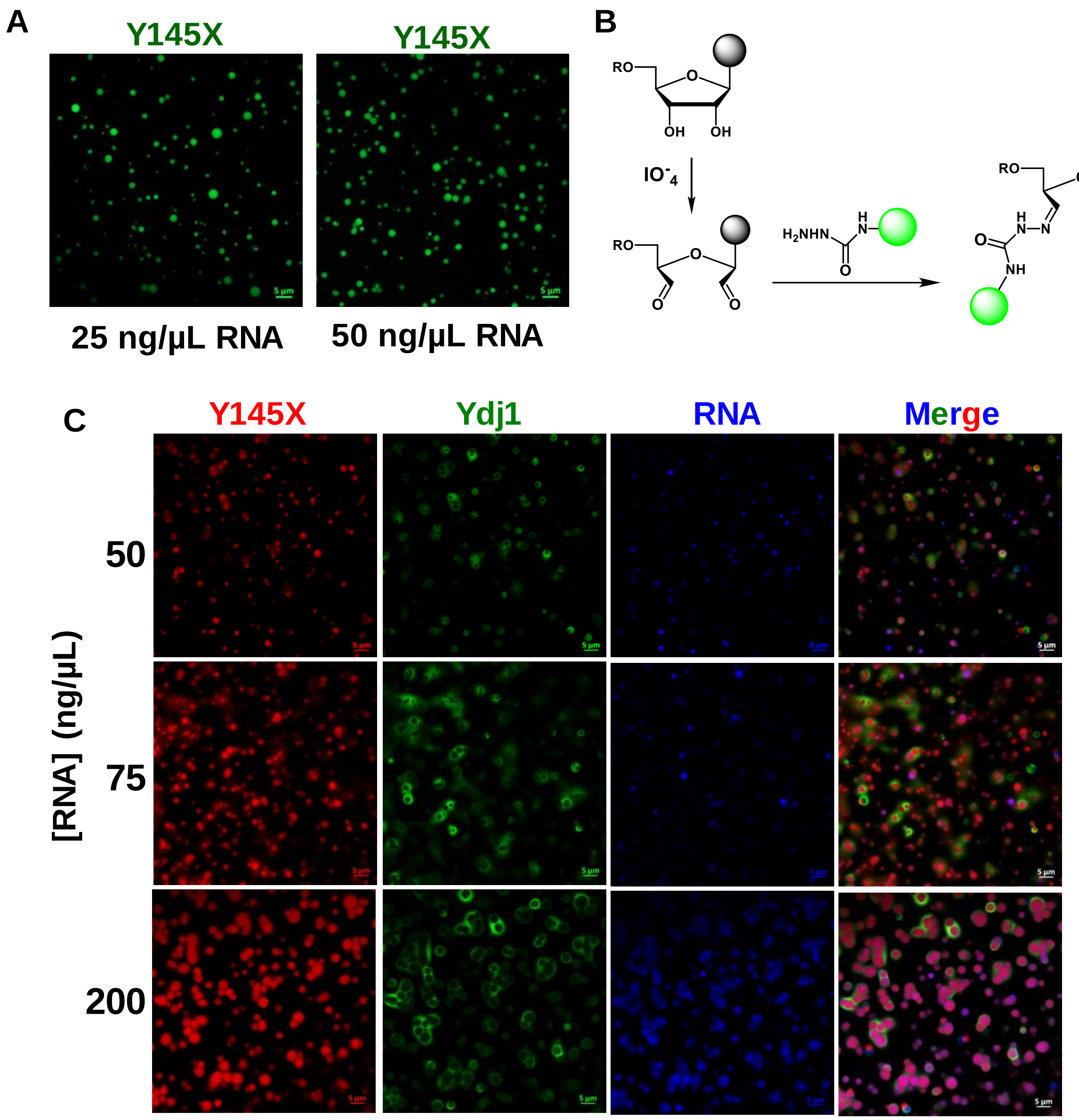


**Fig. S4.** (A) Confocal images of Y145X (AlexaFluor488-labeled) droplets formed in the presence of 25 ng/μl and 50 ng/μl of PolyU RNA. (B) Schematic illustration of RNA labeling with the AlexaFluor488-Hydrazide using periodate chemistry (drawn using ChemDraw). (C) Three-color Airyscan imaging of complex coacervates of Y145X (AlexaFluor647-labeled, red)-RNA (AlexaFluor488-Hydrazide-labeled, blue)-Ydj1 (AlexaFluor594-labeled, green) with increasing RNA concentration. Y145X (100 μM) and Ydj1 (10 μM) concentrations were kept constant. Red, green, and blue colors are used for representation.
